# Modulation of Oncogenic KRAS Signaling by Branched Actin-driven Cell Membrane Protrusions

**DOI:** 10.64898/2026.04.09.717047

**Authors:** Gabriel Muhire Gihana, Kushal Bhatt, Bo-Jui Chang, Vasanth Murali, Felix Zhou, Jungsik Noh, Roshan Ravishankar, Hazel Borges, Jinlong Lin, Jessica Oceguera, Pedro Augusto Nogueira, Lizbeth Perez Castro, Niranjan Venkateswaran, Bingying Chen, Reto Fiolka, Maralice Conacci-Sorrell, Kevin Dean, Gaudenz Danuser

## Abstract

For over three decades, we have known that oncogenic RAS alters the actin cytoskeleton organization and cell surface morphology^1,2^. RAS activates the GTPase RAC1, which triggers the growth of branched actin networks to promote cell membrane protrusions^3,4^. In melanoma, the hyperactive RAC1 mutant, Rac1^P29S^, was recently shown to drive extended lamellipodia, which then empower cell proliferation through sequestration and localized inhibition of the merlin tumor suppressor^5^. This discovery illustrates cell morphological programs not only as outputs but also as regulators of human oncogenic signals. Hence, we wondered whether the pronounced branched actin-driven membrane protrusions (BAMPs) downstream of oncogenic RAS are not mere outputs of RAS signaling but rather an active component in mediating the oncogenic penetrance of RAS mutants. We used volumetric light sheet microscopy and biochemical approaches to investigate the role of BAMPs in regulating the molecular signaling of oncogenic KRAS in pancreatic and lung cancer models. We found that elevated BAMP formation regulated the interaction of oncogenic KRAS with downstream effectors, specifically with the RAC1 GEF TIAM1. This implies that BAMPs amplify their own upstream regulators in a positive feedback. This meritorious cycle upregulates cyclin D1 expression by inactivating the merlin tumor suppressor, independently of the mitogen activated protein kinase pathway (MAPK). In the absence of BAMPs, cells carrying oncogenic KRAS mutations are unable to attain their full penetrance in proliferation. Overall, this work unveils the long-overlooked role of branched actin-driven cell morphology in the functionalization of KRAS mutants as potent oncogenes.

## Introduction

The RAS small GTPases are the most frequently activated oncogenes in human cancer, with more than 20% of patients carrying one or more RAS mutations^6^. Humans have three RAS paralogs: HRAS, KRAS, and NRAS, with KRAS being the most prevalent and involved in deadly pancreatic, lung, and colorectal cancers^6–13^. In 1986, Bar-Sagi and Feramisco showed that oncogenic RAS induces thin cell membrane protrusions known as ruffles and lamellipodia^2^, but ever since, whether this morphological change is only a product of oncogenic RAS or a significant regulator of RAS’ tumorigenic signaling has remained unknown. RAS activates the RAC1 GTPase, and the latter induces ARP2/3-driven cortical branched actin meshes, which push the cell membrane outward to form ruffles and lamellipodia^3,4^. Only recently, data emerged that suggested these morphological programs might participate in the functionalization of RAS mutants themselves, or mutants abnormally activating downstream targets of RAS signals, as powerful oncogenes. In BRAF^V600E^-driven melanoma carrying the hyperactive RAC1^P29S^ mutant, extended lamellipodia formation confer resistance to targeted BRAF/MAPK inhibitor combinations by sequestering and deactivating the Merlin tumor suppressor at the cell periphery^5^. Since Bar-Sagi’s seminal work, it was established that RAC1 and its GEF, TIAM1, are essential for RAS tumorigenesis^14–19^, but how RAC1 mediates the oncogenicity of RAS has remained unknown.

Inspired by the illustration of RAC1’s morphological impact on proliferation signaling^5,20^, we wondered whether the role of RAC1 in RAS cancers could also be related to branched actin formation and subsequent morphological organization of RAS signal transduction. Addressing this question has been made possible by quantitative volumetric imaging of dynamic subcellular structures using high-resolution light-sheet microscopy^21–23^, which provides the sensitivity necessary to detect the effects of morphological shifts on the spatiotemporal organization of molecules at the micron and second scales. In this work, we used pancreatic and lung carcinoma cell line models to investigate how branched actin-driven plasma membrane protrusions (BAMPs), which are RAC1-dependent, affect local concentration of oncogenic KRAS and its interaction with downstream effectors that drive cell proliferation. To ease the discussion, we use the term BAMPs to collectively refer to sheet-like, f-actin, and ARP3-rich cell membrane protrusions, such as ruffles and lamellipodia.

## Results

### Branched actin membrane protrusions (BAMPs) enable local enrichment of oncogenic KRAS

To test the potential role of BAMPs in regulating oncogenic KRAS signaling, we used human cell line models in which all KRAS alleles have oncogenic mutations at codon 12. The SU.86.86 cell line was isolated from a pancreatic adenocarcinoma harboring the KRAS G12D mutation^24,25^. The COR-L23 cell line has the KRAS G12V mutation, and it was isolated from a lung carcinoma^24,26^. Confirmatory DNA sequencing revealed that only mutant KRAS alleles were present in these cell lines, with the indicated respective KRAS mutations (see supplemental file 1). Similar to the fibroblast images shown by Bar-Sagi and Fermisco^1^, SU.86.86 cells display extended, sustained BAMPs. We thus stably expressed ARP3-EGFP as a reporter of branched actin and fluorescently-tagged F-tractin as a reporter of filamentous actin (F-actin)^27^. SU.86.86 cells form ARP3-rich BAMPs in 2D cultures (Fig. 1a, Supplementary Video 1) and in 3D collagen cultures (Fig. 1b, 1c, Supplementary Video 2). As shown by an increased-field-of-view volumetric light-sheet microscope^23^, BAMPs remained robustly present for weeks in SU.86.86 spheroids cultured in 3D collagen matrices (Fig. 1d and Supplementary Video 3). In spheroids, BAMPs formed on the cortex of the outer cells. Using cleared-tissue light-sheet microscopy^22^, we also observed BAMPs on SU.86.86 cells in mouse subcutaneous xenografts (Fig. 1e, Supplementary Video 4). In the xenografts, BAMPs appeared morphologically distinct from other plasma membrane protrusions such as filopodia or blebs, resembling BAMPs in 3D collagen cultures (compare Fig. 1e to Fig. 1b).

**Figure 1:**
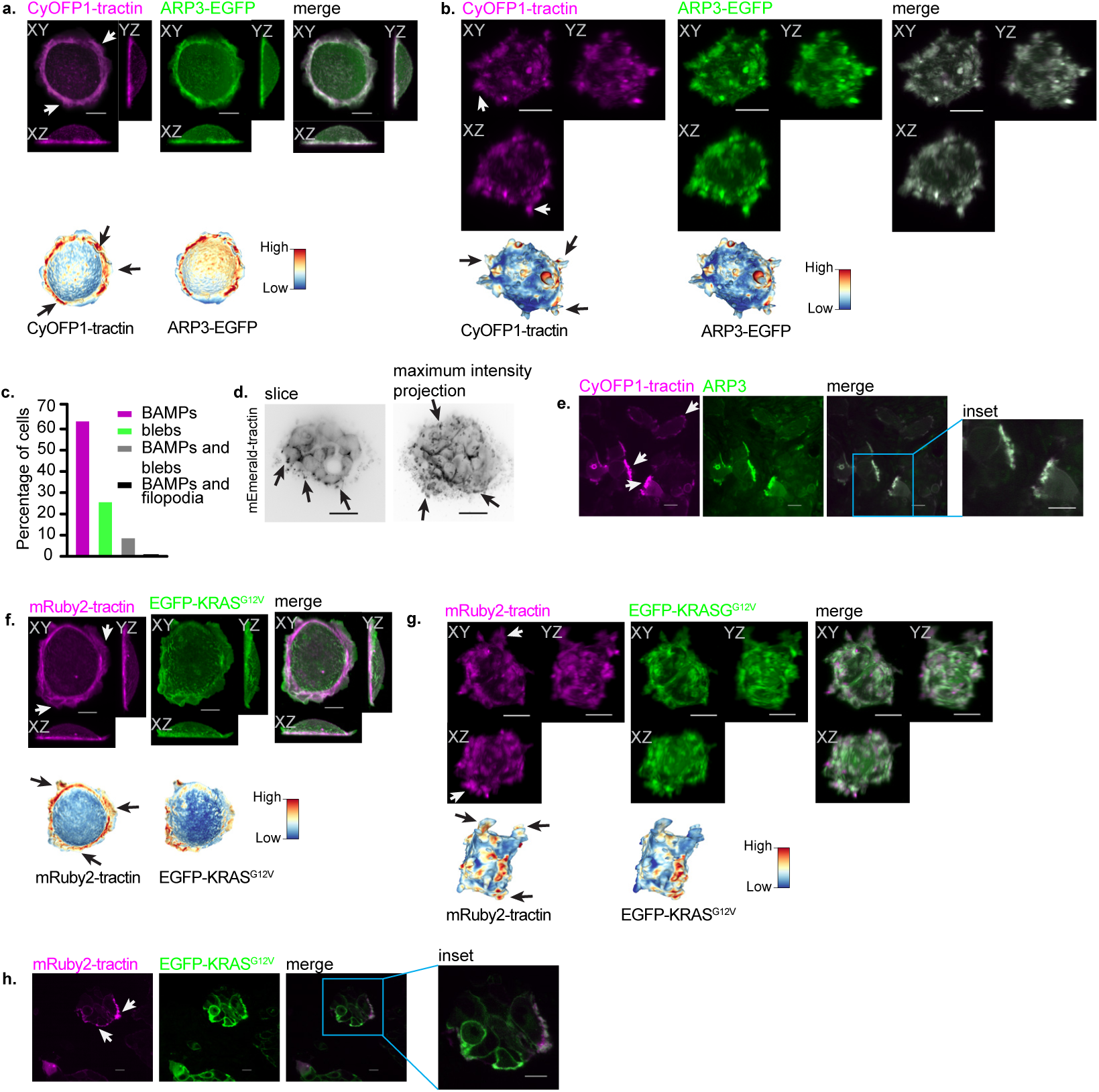
Characterization of branched actin membrane protrusions (BAMPs) in diverse environments. **(a, b)** Representative maximum intensity projection (MIP) images of SU.86.86 pancreatic adenocarcinoma cells plated overnight on a fibronectin-coated coverslip (a) or in 3D collagen matrix (b). Cells were imaged live using a gaussian light sheet on the Field Synthesis microscope21. Heatmaps show fluorescence signals of the indicated fluorophores mapped on the segmented cell surface. Arrows in these and following panels indicate examples of BAMPs extending from the cortical surface of cells. Scale bar: 10µm. **(c)** Frequency of plasma membrane protrusion types formed by SU.86.86 cells cultured in 3D collagen matrix (N=145 cells pooled from three independent replicates). **(d)** Spheroids formed by SU.86.86 cells cultured in 3D collagen for two weeks. **(e)** SU.86.86 cells within a section of a subcutaneous mouse xenograft tumor. The images show one Z plane selected from a volume acquired with a cleared-tissue axially swept light sheet microscope (ct-ASLM)^22^. Scale bar: 10µm. **(f, g)** Representative MIP images of SU.86.86 cells plated overnight on a fibronectin-coated coverslip (f) or in 3D collagen matrix (g). Cells were imaged as described in (a) and (b). Heatmaps show fluorescence signals of the indicated fluorophores mapped on the segmented cell surface. Scale bar: 10µm. **(h)** SU.86.86 cells within a section of a tissue from a subcutaneous mouse xenograft tumor. The images show one Z plane selected from a Z stack acquired on ct-ASLM. Scale bar: 10µm.

We also observed BAMPs in lung cancer COR-L23 cells in 2D culture (Extended Data Fig. 1a, Supplementary Video 5), 3D collagen culture (Extended Data Fig.1b, c, Supplementary Video 6), and in mouse subcutaneous xenografts (Supplementary Video 7). In 3D collagen, more than 70% of both SU.86.86 and COR-L23 cell types formed BAMPs (Figs. 1c and S1c). In addition, supplementary videos 4 and 7 show examples of BAMP formation in many representative SU.86.86 and COR-L23 cells in mouse xenografts, respectively. Therefore, the formation of BAMPs in physiologically relevant contexts is a reproducible and cell-type independent phenotype. Notably, an EGFP-tagged KRAS^G12V^ mutant of the KRAS4B isoform co-localizes with F-actin in the BAMPs of SU.86.86 cells in 2D culture (Fig. 1f), in 3D collagen culture (Fig. 1g), as well as in mouse subcutaneous xenografts (Fig. 1h). The localization of KRAS in BAMPs in physiologically relevant microenvironments suggests that BAMPs may play a role in regulating KRAS’ oncogenic signaling.

### BAMPs amplify the interaction of KRAS mutants with downstream effectors

To investigate if and how BAMPs may regulate the signaling of oncogenic KRAS, we first tested if the oncoprotein was specifically enriched in BAMPs. To this end, we generated BAMPs in a spatiotemporally controlled manner by optogenetic recruitment of the GEF domain of TIAM1 to the cell membrane. This recruitment, based on the iLID system, resulted in local activation of RAC1 and formation of BAMPs^28^. We employed the optogenetic system in MV3 melanoma cells stably expressing ectopic SNAP-tag-KRAS^G12V^ and EGFP fused to a CAAX motif as a membrane marker, which is the same motif that targets KRAS to the cell membrane. We chose the MV3 cells because, in our hands, they reproducibly formed more extended BAMPs compared to other cell lines we tested, making the obtained movies amenable to downstream computational analysis. We observed the recruitment of SNAP-tag-KRAS^G12V^ in light-induced BAMPs (Fig. 2a) and found that the SNAP-tag-KRAS^G12V^ fluorescence intensity, normalized to the intensity of EGFP-CAAX, was significantly higher in BAMPs compared to non-protrusive cell regions (Fig. 2b). Because of the normalization of SNAP-tag-KRASG12V fluorescence intensity to EGFP-CAAX intensity, this result is not merely an artifact of limited optical resolution of membrane folds but indicates bona fide enrichment of oncogenic KRAS in BAMPs.

**Figure 2:**
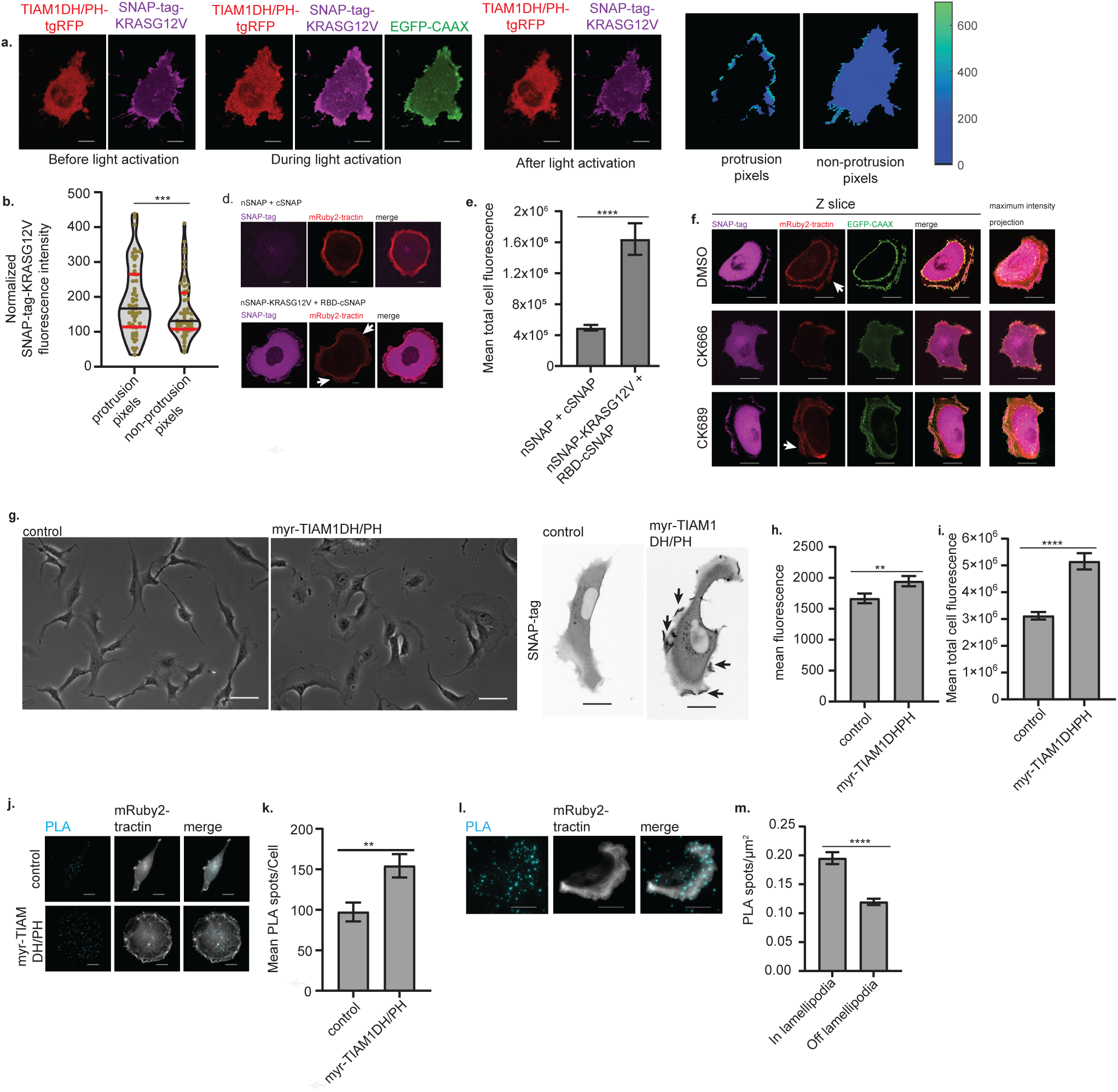
BAMPs concentrate oncogenic KRAS locally and amplify its interactions with downstream effectors. **(a)** Representative images of an MV3 cell expressing Tiam1DH/PH-tgRFPt-SSPBR73Q, myristoylated mTurquoise2-iLID, SNAP-tag-KRASG12V, and EGFPCAAX. The cells were cultured on fibronectin-coated coverslips overnight and imaged with a spinning disk confocal microscope. The heatmaps show the fluorescence intensity of SNAP-tag-KRASG12V normalized to the EGFP-CAAX intensity. Scale bar: 10μm. **(b)** SNAP-tag-KRASG12V fluorescence intensity normalized over EGFP-CAAX intensity. Dots: per-cell mean values of normalized intensity for pixels located in protrusive versus non-protrusive cell regions; horizontal lines: quartiles, with the black line showing the median (n=53 cells pooled from three independent replicates. ***: p value = 0.0002, Wilcoxon matched-pairs signed rank test, two-tailed). **(c)** Top: MIP images of an SU.86.86 cell expressing the N terminal fragment (nSNAP) and C terminal fragment (cSNAP) of the Split SNAP-tag. Bottom: MIP images of an SU.86.86 cell expressing nSNAP-KRASG12V and RBD-cSNAP fragments of the Split SNAP-tag. RBD: RAS Binding Domain of c-RAF1. The cells were imaged on a spinning disk confocal microscope. Scale bar: 10μm **(d)** Per-cell mean of the Z-integrated total Split SNAP-tag fluorescence of SU.86.86 cells represented in (c). Plots show the mean with error bars showing the standard error of the mean (SEM) of n=60 pooled from 3 independent replicates; ****: p value < 0.0001. **(e)** MIPs of of SU.86.86 cells expressing nSNAP-KRASG12V and RBD-cSNAP. The cells were treated with DMSO (0.2%), CK666(200μM), or CK689 (200μM). Scale bar: 10μm **(f)** HPNE cells expressing a doxycycline-inducible TIAM1 DH/PH GEF domain, fused to the myristoylation domain of Lyn11 for cell membrane targeting (myr-TIAM1 DH/PH). The cells also expressed nSNAP-KRASG12V and RBD-cSNAP fragments of the Split SNAP-tag. Brightfield (left) and MIPs (right) are shown. The cells indicated as myr-TIAM1DH/PH were exposed to doxycycline (2μg/mL), while control cells were exposed to the solvent (water). Scale bar: 10μm. **(g, h)** Mean (g) and total (h) fluorescence intensity of the cells represented in (f), measured on sum intensity projections of Z stacks. Plots show the mean with SEM error bars (n = 286 cells pooled from three independent replicates; ****: p value < 0.0001). **(i)** Proximity ligation assay (PLA) detecting the interaction of TIAM1 and KRASG12V in MIA PaCa-2 cells expressing a doxycycline-inducible myr-TIAM1 DH/PH. Images were acquired in a widefield mode on a Delta Vision OMX microscope system. Control cells were not exposed to doxycycline. Scale bar: 10μm. **(j)** Mean number of PLA spots/cell counted in the cells represented in (i). The plot shows the mean with SEM error bars (n=12 cells from one replicate; **: p value = 0.0058). **(k)** PLA for TIAM1-KRASG12D interaction in SU.86.86 cells. Images were acquired in a TIRF mode on a Delta Vision OMX microscope system. Scale bar: 10μm. **(l)** PLA spot density in lamellipodia vs other parts of the ventral cortex calculated in the cells represented in (k). The plot shows the mean with SEM error bars (n = 23 cells from one replicate; ****: p value < 0.0001). For d, g, h, j, l: P values were calculated using a two-tailed Mann-Whitney test. Arrows indicate examples of BAMPs formed on the cortex of the cells.

We reasoned that the enrichment of oncogenic KRAS in BAMPs might enhance the interaction between KRAS and its downstream effectors. We probed the level and the spatial organization of KRAS-effector interactions using a split SNAP-tag reporter^29^. To engineer the reporter, we fused the N-terminal fragment of the SNAP-tag protein (nSNAP; amino acids 1-91) to oncogenic KRAS^G12V^, while the C-terminal fragment (cSNAP; amino acids 92-182) was fused to the RAS binding domain (RBD) of the c-RAF1 effector (amino acids 51-131). We expected higher SNAP-tag fluorescence intensity recovery when KRAS^G12V^ and RBD are proximal enough to reconstitute the SNAP-tag protein. Indeed, compared to the cells expressing the nSNAP and cSNAP fragments only, cells expressing nSNAP-KRAS^G12V^ and RBD-cSNAP yielded four times higher fluorescence intensity (Fig. 2c, d), implying that total cellular fluorescence recovery of the split SNAP-tag can be used to probe the level of oncogenic KRAS-effector complex formation.

The fluorescence of the split SNAP-tag was strongest in BAMPs and the perinuclear region of SU.86.86 cells. However, when we treated the cells with CK-666, a small molecule inhibitor of branched actin^30^, the BAMPs disappeared, followed by spatial reorganization of the split SNAP-tag signal (Fig.2e). Neither the DMSO solvent alone nor the inactive control compound, CK-689, altered BAMP formation or split SNAP-tag fluorescence spatial organization (Fig. 2e). These results suggest that, as cells form BAMPs, they gain microdomains in which oncogenic KRAS interacts with downstream effectors.

To test if we could induce KRAS-effector complex formation at the cellular level by acute formation of BAMPs, we probed KRAS^G12V^-RBD interaction using the split SNAP-tag reporter in HPNE human pancreatic duct cells. These cells undergo multiple passages and retain an elongated morphology in 2D culture even in the presence of ectopic oncogenic KRAS, making the effect of BAMP induction readily appreciable. In these cells, we induced BAMPs by engineering and expressing the TIAM1 GEF domain (amino acids 1030-1406), to which we fused the membrane targeting myristoylation domain of the Lyn11 kinase (amino acids 1-11), and a 3X FLAG tag. The myristoylated TIAM1 GEF domain (myr-TIAM1DH/PH) was placed under a doxycycline-inducible promoter. The myr-TIAM1DH/PH, which was readily expressed in presence of doxycycline (Extended Data Fig. 2a), triggered BAMP formation and yielded both significantly higher mean and total SNAP-tag fluorescence intensity (Fig. 2f, g, h). The higher mean fluorescence intensity is particularly notable, given that myr-TIAM1DH/PH led to an increased cell size (Extended Data Fig. 2b, Fig. 2f). Nonetheless, the higher total cell fluorescence intensity was not simply an effect of the increased cell size, because cells expressing myr-TIAM1DH/PH had significantly lower total fluorescence intensity in the absence of KRAS^G12V^ and RBD (Extended Data Fig. 2c). We also confirmed that doxycycline alone did not alter cell shape (Fig. 2Sd), neither did it increase cell size or the fluorescence intensity (Extended Data Fig. 2e, f, g). In addition, doxycycline did not alter the expression level of nSNAP-KRAS^G12V^ or RBD-cSNAP (Extended Data Fig. 2h). Together, these results indicate that BAMPs enhance the formation of oncogenic KRAS-effector complexes.

TIAM1 plays a central role in RAS-induced branched actin formation and cellular morphogenesis, because it links RAS to RAC1 by acting as both a RAS effector and a RAC1 GEF^3,31^. Therefore, we investigated the effect of BAMPs on oncogenic KRAS-TIAM1 interactions. To this end, we used a proximity ligation assay (PLA) in patient-derived MIA-PaCa2 cells. Despite having both KRAS alleles mutated to KRAS G12C^24,32^ (Supplemental file 1), these human pancreatic carcinoma cells do not form prevalent BAMPs in 2D culture, providing an adequate study system to induce BAMPs and interrogate their effect on oncogenic KRAS^G12C^-TIAM1 interactions. Upon expression of myr-TIAM1DH/PH, MIA-PaCa2 cells spread out and formed extended BAMPs, which led to increased oncogenic KRAS^G12C^-TIAM1 interactions, quantified as the total number of PLA spots in the cell (Fig. 2i, j). Given the elevated interaction between oncogenic KRAS and the c-RAF1 RBD observed in BAMPs with the split SNAP-tag system, we wondered whether the formation of KRAS-TIAM1 signaling complexes would be promoted also in a system with baseline BAMPs. We performed a PLA between KRAS^G12D^ and TIAM1 in SU.86.86 cells. Indeed, we observed a higher density of PLA spots in lamellipodia compared to the rest of the cell (Fig. 2k, l). Thus, we conclude that BAMPs provide more generally a microdomain that enhances oncogenic KRAS’ interaction with its downstream effectors, including TIAM1.

### BAMPs promote cyclin D1 expression and cell proliferation downstream of oncogenic KRAS, independently of MAPK

Given BAMPs’ function in organizing microdomains of KRAS signal transduction, we wondered whether this morphogenic program more broadly promotes the penetrance of KRAS mutants as oncogenes. RAS mutants trigger multiple signaling pathways, principally the mitogen activated protein kinase (MAPK) pathway, which promotes cell proliferation^33^. Proliferative signals promote the expression of cyclins and the activation of cyclin dependent kinases, which drive the cell cycle^34,35^. As expected, in SU.86.86 cells, inhibition of KRAS G12D with the pan-KRAS inhibitor BI-2865^36^ led to a significant decrease in cell proliferation and MAPK signaling, as indicated by the reduction of ERK1/2 phosphorylation (Extended Data Fig. 3A, B). Treatment of SU.86.86 cells with BI-2865 also abrogated BAMPs, which was morphologically compensated by bleb formation (Extended Data Fig. 3C, D). This result shows that, in this cell line, oncogenic KRAS is responsible for BAMP formation, making it an ideal model to investigate the role of oncogenic RAS-driven BAMPs in RAS signaling.

To assess the penetrance of BAMP inhibition at the cell population level, we developed a computational analysis pipeline to quantify the rate of BAMP formation (See Online Methods). As qualitatively shown before (Fig. 2e), this pipeline now confirmed that inhibition of Arp2/3 activity by CK-666 disrupted BAMP formation significantly (Fig. 3a, b), which was accompanied by a striking reduction in cell proliferation (Fig. 3c). Both BAMP disruption and reduction of cell proliferation under CK-666 but not under CK-689 treatment were reproduced in 3D culture in collagen matrices (Fig. 3d, e). The CK-666-triggered reduction in cell proliferation correlated with a decrease in cyclin D1 expression (Fig. 3f, g); but surprisingly, ERK1/2 phosphorylation was increased (Fig. 3f, h). We obtained similar results in COR-L23 cells treated with CK-666 (Extended Data Fig. 4), and in SU.86.86 cells treated with wheat germ agglutinin (WGA), a lectin that limits cell membrane protrusions by binding to the glycocalyx^37,38^ (Fig. 3i, j, k, l, m). These results imply that, despite being constitutively active, oncogenic KRAS’ ability to boost cyclin D1 expression and cell proliferation depends on BAMPs, at least in the contexts where cells form these protrusions. Additionally, the results suggest that BAMP-dependent upregulation of cyclin D1 expression and cell proliferation downstream of oncogenic KRAS is independent of MAPK. In support of this observation, inhibition of MAPK alone yields only a partial decrease in cell proliferation and cyclin D1 levels, comparable to the loss caused by BAMP inhibition. Co-inhibition of BAMPs and MAPK leads to near-total abolishment of cyclin D1 proteins and an additive decrease in cell proliferation (Fig. 3n, o, p). Hence, in presence of oncogenic KRAS, BAMPs sustain cyclin D1 expression and cell proliferation even when MAPK is downregulated.

**Figure 3:**
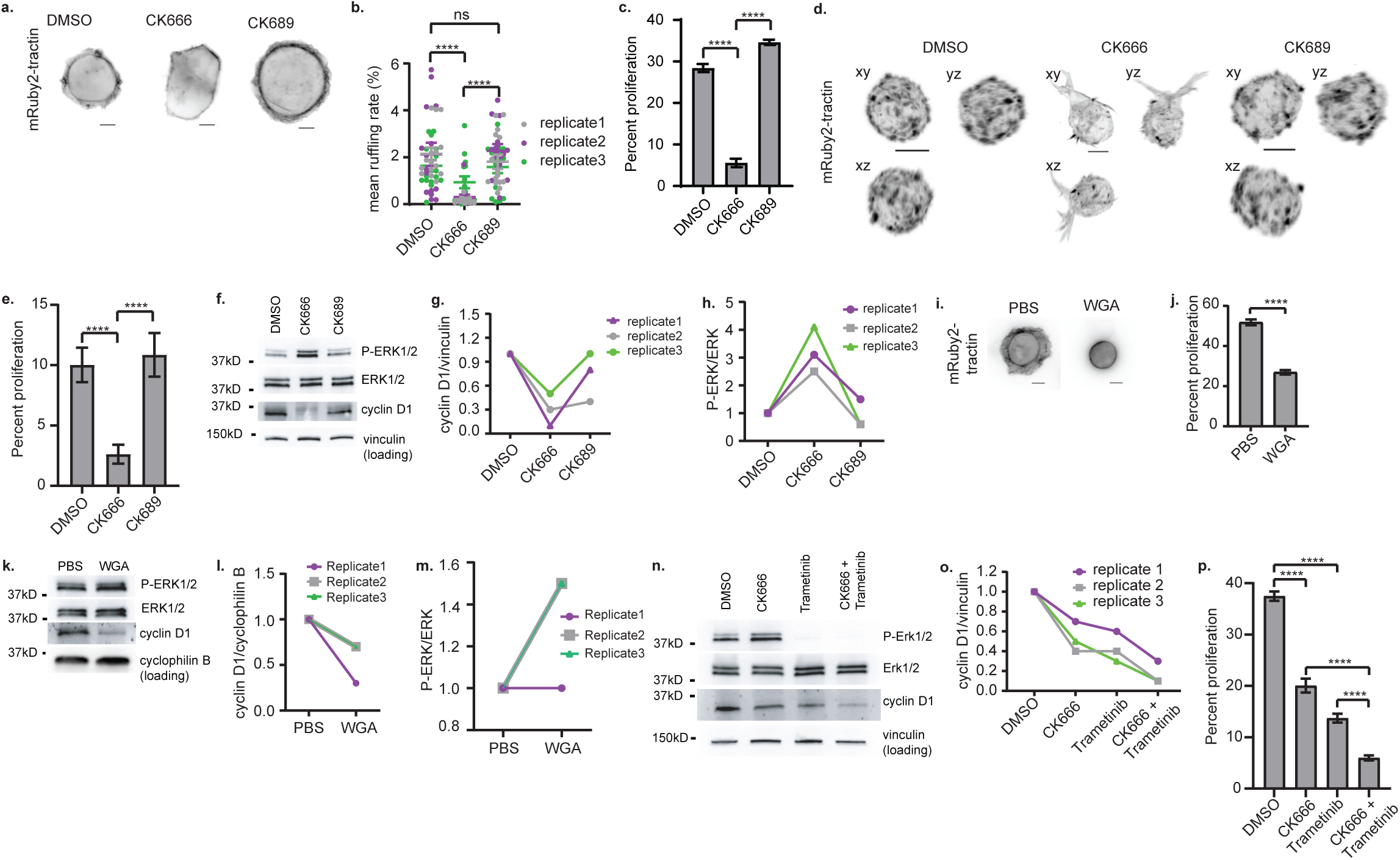
BAMPs promote cyclin D1 expression and cell proliferation downstream of oncogenic KRAS, independently of MAPK. **(a)** Epifluorescence images of SU.86.86 cells treated with DMSO, CK666, or CK689. Scale bar: 10µm. **(b)** Mean membrane ruffling rate computed in the cells represented in (a). Dots represent individual image frames; error bars represent SEM (n = 45 image frames pooled from three independent replicates, with each image containing 1-10 cells; ****: p value <0.0001). **(c)** Proliferation rate of SU.86.86 cells treated with DMSO, CK666, or CK689. The plot shows the mean with SEM error bars (n = 45 images analyzed from three independent replicates; ****: p value <0.0001). **(d)** MIP images of SU.86.86 cells cultured in 3D collagen matrices and treated with DMSO, CK666, or CK689. Cells were imaged using a gaussian light sheet on the Field Synthesis microscope22. Scale bar: 10µm. **(e)** Proliferation rate of SU.86.86 cells treated with DMSO, CK666, or CK689 in 3D collagen culture. The plot shows the mean with SEM error bars (n = 30 images pooled from three independent replicates; ****: p value <0.0001). **(f)** Western blotting for phosphorylated ERK1/2 and cyclin D1 in SU.86.86 cells treated DMSO, CK666, or CK689. **(g, h)** Quantification of band intensity on the western blot images represented in (f). **(i)** MIP images of SU.86.86 cells cultured in PBS (solvent control) or wheat germ agglutinin (WGA; 10µg/mL). Scale bar: 10µm. **(j)** Proliferation rate of SU.86.86 cells treated with PBS or WGA. The plot shows the mean with SEM error bars (n = 30 image frames from three independent replicates; ****: p value <0.0001). **(k)** Western blotting for phosphorylated ERK1/2 and cyclin D1 in SU.86.86 treated with PBS or WGA. **(l, m)** Quantification of band intensity on the western blot images represented in (k). **(n)** Western blotting for phosphorylated ERK1/2 and cyclin D1 in SU.86.86 cells treated with DMSO, CK666, trametinib, or CK666+trametinib. **(o)** Quantification of band intensity on the western blot images represented in (n). **(p)** Proliferation rate of SU.86.86 cells treated with DMSO, CK666, CK689 or CK666 +trametinib. The plot shows the mean with SEM error bars (n = 73 images pooled from three independent replicates; ****: p value <0.0001). DMSO was used at 0.2%, CK666 and CK689 at 200µM, and trametinib at 10nM. The treatment was for 48 hours. Vinculin and cyclophilin B were used for loading control in western blots. All P values were calculated using a two-tailed Man-Whitney test. For c, e, j, and p, each analyzed image contained at least dozens of cells.

### BAMPs mediate merlin inhibition downstream of oncogenic KRAS

The merlin tumor suppressor inhibits cyclin D1 expression, and it counteracts RAS tumorigenesis^39–42^. We recently found that hyperactive RAC1^P29S^, but not wildtype RAC1, induces extended lamellipodia that sequester merlin and enhance inactivating phosphorylation by the PAK1 kinase^5^. This led us to hypothesize that the BAMPs driven by constitutively active KRAS would be robust enough to promote the inhibitory phosphorylation of merlin even downstream of wildtype RAC1. Indeed, BAMP inhibition with CK-666 reduced merlin phosphorylation in SU.86.86 (Fig. 4a, b) and COR-L23 (Fig. 4c, d) cells. Treating SU.86.86 cells with WGA yielded similar results (Fig. 4e, f), indicating that this phenotype is not cell-type specific and is independent of the method used to perturb BAMPs. In line with the involvement of BAMPs in regulating merlin, endogenously tagged merlin localizes in BAMPs (Fig. 4g).

**Figure 4:**
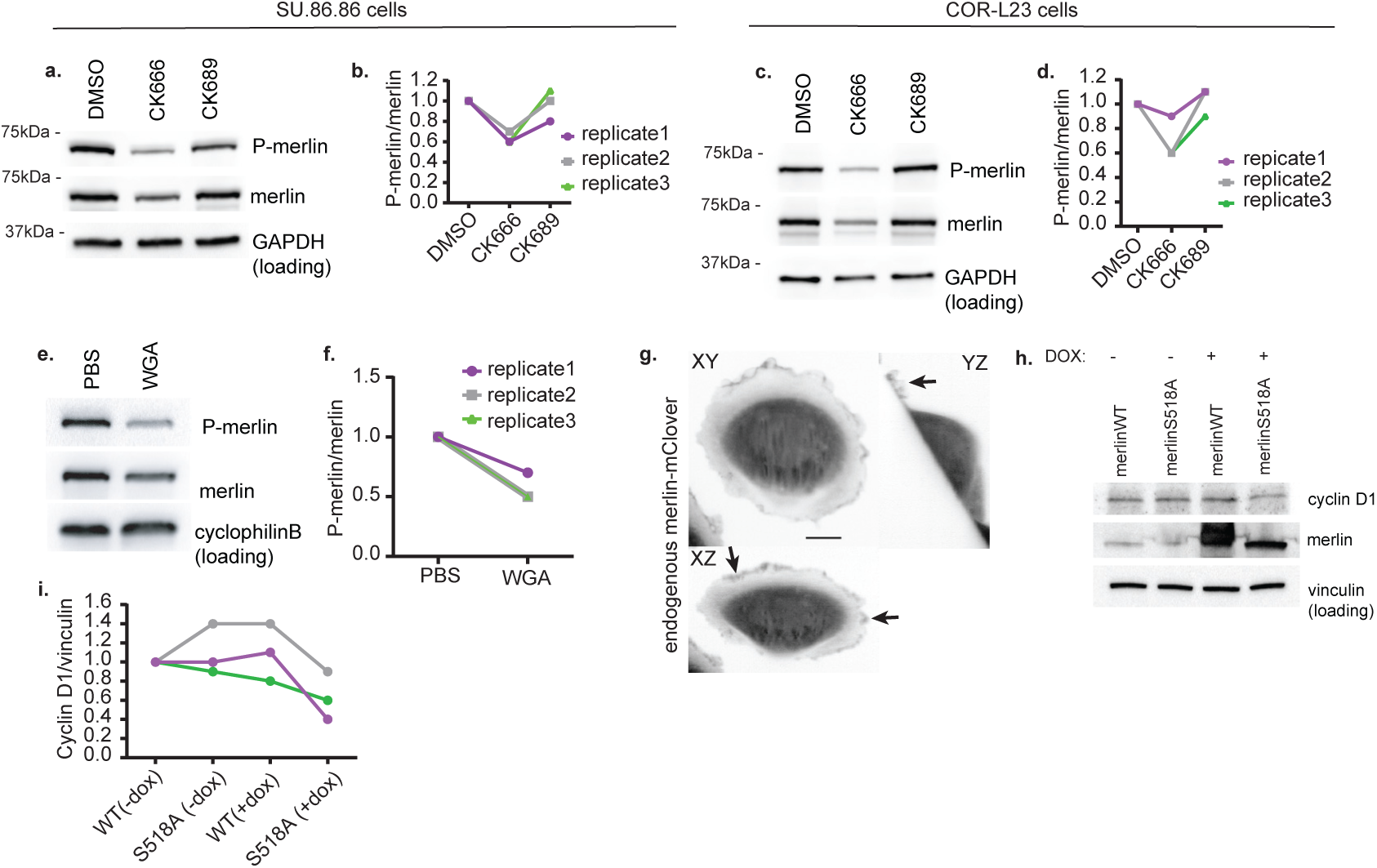
BAMPs promote merlin inhibition downstream of oncogenic KRAS. **(a)** Western blotting for phosphorylated merlin in SU.86.86 cells treated with DMSO, CK666, or CK689. GAPDH was used for loading control. **(b)** Quantification of band intensity on the western blot images represented in (a). **(c)** Western blotting for phosphorylated merlin in COR-L23 cells treated with DMSO, CK666, or CK689. GAPDH was used for loading control. **(d)** Quantification of band intensity on the western blot images represented in (c). **(e)** Western blotting for phosphorylated merlin in SU.86.86 cells treated with PBS (solvent control) or wheat germ agglutinin (WGA). Cyclophilin B was used for loading control. **(f)** Quantification of band intensity on the western blot images represented in (e). **(g)** Maximum intensity projection images of a COR-L23 cell expressing endogenously tagged merlin-mClover. The cells were plated on fibronectin-coated coverslip overnight and imaged using a gaussian light sheet on the Field Synthesis microscope22. Scale bar: 10µm. Arrows indicate examples of merlin localized in BAMPs. **(h)** Western blotting for cyclin D1 and merlin in SU.86.86 with merlin wildtype (WT) or the S518A merlin phosphomutant under a doxycycline (DOX)-inducible promotor. Vinculin was used for loading control. DMSO was used at 0.2%, CK666 and CK689 at 200µM, and WGA at 10µg/mL. **(i)** Quantification of band intensity on the western blot images represented in (h). Doxycycline (dox) was used at 2µg/mL).

We reasoned that, if the diminution of cyclin D1 levels upon BAMP inhibition were a result of less phosphorylated and, therefore, more active merlin, the expression of a phosphomutant merlin would restrict cyclin D1 expression comparably to the effect of BAMP inhibition. The inhibitory phosphorylation of Merlin targets serine 518^43–45^. Therefore, we assessed the levels of cyclin D1 expression in SU.86.86 cells ectopically expressing a doxycycline inducible wildtype merlin or S518A phosphomutant. Indeed, overexpression of the S518A phosphomutant, like BAMP inhibition, led to a partial decrease in cyclin D1 levels in all our experimental replicates (Fig. 4h, i). Thus, active and diffusive merlin is sufficient to decrease cyclin D1 expression in the presence of constitutively active KRAS. Overall, this work establishes that oncogenic KRAS induces branched actin-driven membrane protrusions, which via enhanced TIAM1-RAC1-PAK1 signaling, mediate the inhibition of the merlin tumor suppressor, resulting in higher cyclin D1 expression and cell proliferation.

## Discussion

Pioneer studies in the early 1980’s identified RAS human oncogenes as major drivers of cell proliferation and survival^46,47^. In 1986, Bar-Sagi and Feramisco uncovered another effect of oncogenic RAS: the alteration of cell surface morphology by inducing pronounced membrane ruffles and lamellipodia^1^. Ever since, a very significant effort has been invested in studying signaling pathways like the MAPK and PI3K pathways, which drive cell proliferation and survival downstream of oncogenic RAS^47–62^. However, much less attention, if any, has been given to the potential role of cell morphogenesis in regulating molecular signals downstream of oncogenic RAS. Here, we show that RAS-dependent branched actin-driven membrane protrusions (BAMPs) promote the penetrance of oncogenic KRAS mutants by enhancing proliferation signals in human pancreatic and lung cancer cells. We found that the very narrow spaces in BAMPs locally concentrate oncogenic KRAS and enhance its interaction with downstream effectors, specifically TIAM1, which activates Rac1 and thus reinforces BAMP formation in positive feedback.

How could a BAMP locally concentrate RAS? There are several plausible mutually non-exclusive mechanisms that will have to be explored in detailed biophysical and biochemical analyses. In the simplest scenario, the narrow space of BAMPs restricts the diffusion of KRAS and its direct effectors (TIAM1) and effectors further downstream (RAC1), increasing the probability of the molecules to interact and transmit their signaling state. Another possibility could arise from the posttranslational modifications of the RAS proteins^63–65^, some of which, hypothetically, might afford them higher affinity for BAMP microdomains. Alternatively, the dense packing of branched actin may promote scaffolding effects: proteins that bind both branched actin and RAS would, conceivably, lead to a retention of RAS inside the BAMP, which in turn could again promote the frequency of interactions between KRAS molecules and its downstream effectors. Regardless of which exact mechanism of Ras enrichment is at play at the molecular scale, our data shows heightened proximity of oncogenic KRAS and TIAM1. Critically, it also shows that blocking the pathways of branched actin formation downstream of TIAM1-RAC1 signaling or mechanically restricting BAMP expansion reduces cell proliferation. This indicates that BAMPs are not merely an epiphenomenon of enhanced Ras-Rac signaling in Ras-transformed cells but that they play a critical role in empowering Ras mutations as cancer signals.

A striking result of the presented experiments indicates that BAMP-mediated proliferation signaling by KRAS mutations bypasses the canonical MAPK signaling pathway. We found that BAMP inhibition decreased cyclin D1 expression and cell proliferation all the while MAPK was hyperactivated (Fig. 3A-M, Extended Data Fig. 3). When we inhibited MAPK in our cellular models, we observed only a partial decrease in cyclin D1 (Fig. 3N-P) implying that, in cases where KRAS-transformed cancer cells can form BAMPs, these morphogenic structures afford the cells a MAPK-independent proliferative advantage. These data corroborate previous evidence and provide a mechanism for the limited effect of MAPK inhibitors against RAS-addicted cancers^66–68^. Furthermore, in alignment with previous observations in melanoma with amplified lamellipodia extension downstream of a hyper-active RAC1 mutant, we found that KRAS-dependent BAMPs promote MAPK-independent proliferation signals via spatially regulated inhibition of the tumor suppressor Merlin, which is also an inhibitor of BAMP formation^5,69^. Thus, we propose that a significant portion of KRAS’ penetrance as a proliferation-driving oncogene is mediated by a double negative feedback loop.

Where could BAMPs form and be critical for RAS oncogenic penetrance and tumor growth *in-vivo*? Intravital imaging has revealed the formation of lamellipodia on the cortex of mouse and human cancer cells engrafted into mice^70,71^. In this work, we found that BAMPs form at the surface of peripheral cells in subcutaneous mouse xenograft tumors of human pancreatic and lung cancer cells (Figs. 1E, Supplementary Videos 4, 7), and we have shown that oncogenic KRAS localizes in the xenograft BAMPs (Fig. 1H). Compared to intravital imaging, our cleared-tissue light sheet microscope^22^ allowed us to expand the study sample size and to confirm that BAMP formation *in-vivo* is a penetrant and cell-type independent phenomenon. The uniting feature between our study and others is that BAMPs tend to form at the periphery of the tumor mass. Intriguingly, multiple studies exploring human cancers have observed more cell proliferation at the tumor periphery compared to the center of tumors originating from lung, kidney, skin, and soft tissue^72–76^. The correlation between the intratumoral proliferative zones with the zones where BAMPs form support our conjecture that protrusions play a role in regulating cancer cell proliferation *in-vivo*.

Overall, this work reveals that the cellular morphological change induced by oncogenic RAS is not a mere phenotypic manifestation, but rather an important regulator of oncogenic RAS’ interactions with downstream effectors. Importantly, our results show that oncogenic KRAS relies on BAMPs to attain full penetrance despite being persistently active. Recent successful development and FDA approval of oncogenic KRAS inhibitors have raised hopes, but oncogenic RAS remains a significant therapeutic challenge^66^. Indeed, resistance to the new targeted KRAS inhibitors has been reported alreday^77–84^. Our work reveals BAMP formation, a nongenetic process, as an important modulator of RAS oncogenic penetrance. This opens a new window to therapeutic approaches that complement targeted drugs by exploiting cellular processes that configure the biochemical conditions of mutant signaling.

## Materials and methods

### Cell lines and cell culture

SU.86.86 cells (ATCC, CRL-1837) and COR-L23(Millipore Sigma, 92031919) were cultured in RPMI-1640 medium supplemented with 10% fetal bovine serum (FBS) and 1% antibiotic-antimycotic (Thermo Scientific, 15240062). MIA-PaCa2 cells (ATCC, CRL-1420) were cultured in Dulbecco’s Modified Eagle’s Medium (DMEM, Gibco) supplemented with 10% FBS, 2.5% horse serum, and 1% antibiotic-antimycotic. The HPNE cells were a gift from Rolf Brekken (UT Southwestern Medical Center, Dallas TX), and they were cultured in DMEM supplemented with 10% FBS and 1% antibiotic-antimycotic. MV3 cells were a gift from Peter Friedl (MD Anderson Cancer Center, Houston TX), and they were cultured in DMEM supplemented with 10% FBS and 1% antibiotic-antimycotic. All cells were cultured at 37°C and 5% CO^2^ in humidified incubators. All cells used in this study showed no mycoplasma contamination.

For 2D culture, cells were seeded on conventional tissue-culture treated plastic dishes (Corning). For 3D collagen culture, 500 thousand cells were seeded in 1.8mg/mL bovine collagen (Advanced Biomatrix, #5005) supplemented with 0.01M sodium hydroxide in a phosphate buffered saline solution (PBS). The cell-collagen mixture was incubated at 37oC for 1 hour for the collagen to solidify before adding appropriate culture medium. For imaging, cells were incubated overnight. For CK-666, DMSO, and CK-689 treatments, cells were incubated for 48 hours. To make spheroids, cells were incubated for multiple days to multiples weeks, refreshing media every 2-3 days.

### Cloning

For cloning pertaining to CRISPR gene modification, see the CRISPR tagging section below. All other genes cloned in this study were inserted into either the pLVX bicistronic IRES lentiviral vector or the pTRE3G tetON lentiviral vector, using the New England Biolabs Gibson assembly protocol. The pTRE3G empty vector was generated by linearizing pLX-TRE-dCas9-KRAB-MeCP2-BSD (Addgene #140690) with the following primers: fwd: 5’-AAGGGTTCGATCCCTACC-3’, Rev: 5’-GGTGGCAAATTCGAATTCG-3’. The linear PCR product was then circularized and transformed into competent cells using the KLD reaction of the Q5 site directed mutagenesis kit (New England Biolabs, E0554S). The plasmid was sequenced for verification. The genes cloned into the pLVX vector were inserted after digestion of the backbone plasmid with EcoRI and BamHI. For cloning into the pTRE3G tetON expression vector, genes were inserted into the vector linearized with BstBI. Plasmid constructs were verified through whole-plasmid sequencing (Plasmidsaurus). All the plasmids used in this study are listed in Supplementary Table 1.

### Lentivirus infection and ectopic gene expression

For stable expression of mRubyt2-tractin, CyOFP1-tractin, mEmerald-tractin, Arp3-EGFP, EGFP-KRASG12V, SNAP-tag-KRASG12V, nSNAP-KRASG12V, RBD-cSNAP, mTiam1(64-437)- tgRFPt-SSPB R73Q, mTiam1(64-437)-tgRFPt-SSPB R73Q and EGFP-CAAX, we used lentiviral integrating plasmids. Lentiviral particles were produced in human embryonic kidney (HEK) 293T cells by transfecting cells with 5 μg of pMD.2 g 5 μg of psPax2 and 5 μg of the plasmid harboring the gene of interest. A 1:3 DNA:PEI (polyethylenimine) ratio was used for transfection, and the HEK cells were incubated in viral medium overnight. The viral medium was replaced with fresh medium the next day and incubated for 24 hours. For transduction, the viral medium was filtered through a 4.5µm syringe filter, mixed with fresh medium (1:1 ratio) supplemented with polybrene (2 μg/ml), and added to the cells to be transduced. The media was replaced with appropriate culture media for the transduced cells after 24 hours, and antibiotic treatment or flow cytometry sorting was initiated afterwards to select for positively transduced cells.

### CRISPR tagging

We used CRISPR to add m-Clover at the C terminus of NF2, which encodes merlin. Two guides were generated using CRISPOR^85^: gRNA1: 5’-CTTCTTTGAAGAGCTCTAGC-3’ and gRNA2: 5’-TCTTTGAAGAGCTCTAGCAG-3’. The guides were cloned into pSpCas9(BB)-2A-Puro (PX459) V2.0 (Addgene plasmid # 62988). The donor vector was generated using pMA-Tia1L^86^, which was a gift from Tilmann Bürckstümmer (Horizon Genomics, Austria). The donor vector was designed to have the mClover cDNA between 350 bp of NF2 homologous arms each, on either side. The plasmids were transfected into COR-L23 cells using lipofectamine 3000 according to manufacturer’s protocol. 48 hours post-transfection, we sorted for mClover-positive cells using FACS. After sorting, the cells were expanded and validated by sequencing the NF2 c-terminus.

### Cell proliferation assay

The percentage of proliferating cells was determined by capturing the fraction of the cells in S phase of the cell cycle using the ethynyl deoxyuridine (EdU) click chemistry kit for imaging (ThermoFisher Scientific). 40,000 cells were seeded into a well of a twelve-well tissue culture plate and incubated for 48 hours. The cells were then incubated with 10µM EdU for 1 hour, fixed with 4% paraformaldehyde, and permeabilized with 0.5% triton. The incorporated EdU was labeled according to the manufacturer’s protocol, and all nuclei were labeled with DAPI (Invitrogen, D21490). For image acquisition see protocols in the microscopy section below. To determine the percentage of EdU positive cells, we analyzed the acquired images using Cell Profiler as described before^87,88^. Briefly, nuclei were segmented for both the EdU (Cy5 or FITC) and DAPI channels. The same thresholding was applied for samples being compared (DMSO versus CK666 versus CK689 treatments, for example). The total number of positive pixels were determined for each channel, and the ratio of EdU positive pixels over DAPI positive pixels was taken as the rate of cell proliferation, expressed as a percentage.

### Mouse subcutaneous xenograft

Cells were grown to a confluency between 40 and 60% in 15cm plates and detached from the culture plates using trypsin. The cells were washed twice and resuspended in PBS. 2 million cells were injected subcutaneously in both flanks of immune compromised NOD.Cg-Prkdcscid/J mice (Jackson Labs). When the tumors became palpable, they were dissected out, fixed with 4% paraformaldehyde for 24 hours, washed and stored in PBS at 4°C. The tumors were then sliced into 0.5-2mm slices before tissue clearing.

### CUBIC Tissue Clearing

Tumor samples were harvested and fixed overnight in 4% PFA shaking at 4°C. Fixative solution was removed with 3 washes with 0.02% sodium azide PBS, refreshing the PBS every 2 hours at RT. Samples were sliced in 0.5-2mm slices using a tumor matrix (TED PELLA 15020), then slices were immersed in 50% CUBIC-L (TCI Chemicals, T3740) with constant rotation overnight at 37°C. The next day, samples were placed in 100% CUBIC-L with constant rotation at 37°C, refreshing the delipidation reagent until the supernatant turned colorless (∼3 days). When delipidation was completed the 100% CUBIC-L was washed out with 3 washes every 2 hours at RT. To match the refractive index, tumor samples were pretreated with 50% CUBIC-R+ (TCI Chemicals, T3741) with constant rotation overnight at RT and the next day immersed in 100% CUBIC-R+, refreshing every day for 2 days. After RI matching, samples were mounted in ctASLM-v2 with its imaging chamber filled with 100% CUBIC-R+ covered with mineral oil.

### Immunolabeling of Arp3 in cleared tissues

Tumor samples were harvested and fixed overnight in 4% PFA shaking at 4°C. Fixative solution was removed with 3 washes with PBS supplemented with 0.02% sodium azide. PBS was refreshed every 2 hours at RT. Samples were sliced in 0.5-2mm slices using a tumor matrix (TED PELLA, 15020). Then the slices were immersed in 50% CUBIC-L (TCI Chemicals, T3740) with constant rotation overnight at 37°C. The next day, samples were placed in 100% CUBIC-L with constant rotation at 37°C, refreshing the delipidation reagent until the supernatant turned colorless (∼3 days). After delipidation, the 100% CUBIC-L was washed out 3 times with PBS every 2 hours at RT. The samples were permeabilized in blocking buffer (0.5%NP40, 10% DMSO, 5% serum, 0.5% Triton X-100 in 1X PBS) with constant rotation overnight at RT. The next day samples were immunolabelled with ARP3 antibody conjugated with Alexa Fluor 488 (Santa Cruz Biotechnology, sc-48344) diluted at 1:100 in staining buffer (0.5%NP40, 10% DMSO, 5% serum, 0.5% Triton X-100 in 1X PBS) rotating for 7 days at RT. The samples were then washed 3 times with the wash buffer (0.5%NP40, 10% DMSO in 1X PBS) every 2 hours at RT. The samples were placed in 50% CUBIC-R+ for 1 day at RT. To match the refractive index (RI), tumor samples were pretreated with 50% CUBIC-R+ (TCI Chemicals, T3741) with constant rotation overnight at RT, and immersed the next day in 100% CUBIC-R+, refreshing every day for 2 days. After RI matching, samples were mounted in the imaging chamber filled with 100% CUBIC-R+ and covered with mineral oil (Fischer Scientific, BP2629).

### Western blots

For western blots, 1 million cells (control cultures) or 1.5 million (CK666 or WGA cultures) were seeded in 15 cm culture dish and incubated for 48 hours. The cells were lysed on the culture dishes using RIPA buffer (150mM sodium chloride, 0.5% sodium deoxycholate, 0.1% sodium dodecyl sulfate, 1mM ethylenediaminetetraacetic acid, 1% NP-140, 0.01 sodium azide, and 50nM Tris-HCL, pH 7.4). The lysates were cleared by centrifugation at 10,000 rpm for 5 minutes, mixed with Laemmli sample buffer (BioRad, 1610747), and stored at -80°C. Proteins were heated at 90°C for 5-10 minutes, separated using electrophoresis on Any KD precast gels (BioRad), and transferred onto PVDF membranes (ThermoFisher Scientific, 88518). The membranes were blocked in 5% low-fat cow milk in a TBST buffer (150mM sodium chloride, 2.7mM potassium chloride, 0.05% Tween 20, 10mM Tris-HCL, pH 7.4) before primary antibody incubation overnight at 4°C. The primary antibodies were diluted and incubated in the blocking solution. After primary antibody incubation, the membranes were washed 3x5 minutes with TBST at room temperature and incubated in blocking solution containing horseradish peroxidase-conjugated secondary antibodies with gentle shaking for 1 hour at room temperature. After the secondary antibody incubation, the membranes were washed 5x5 minutes at room temperature. All washes and antibody incubations were done with gentle rocking. The membranes were incubated in regular enhanced chemiluminescence substrate (ThermoFisher Scientific #32106) or the SuperSignal substrate (ThermoFisher Scientific, #34096) for 5 minutes and imaged on a Syngene G:Box. The acquisition exposure time was optimized from sample to sample, and only images without any saturated camera pixels were saved and used in downstream analysis.

### Western blot band densitometry

The density of the protein bands on western blot membrane images were determined using the ImageJ Gels analysis tool. To calculate the relative density, the density of all bands was normalized to the density of the bands from the solvent control sample (DMSO for CK-666 treatment or PBS for WGA treatment) and to the density of the bands of the corresponding loading control proteins.

### Antibodies

The following antibodies were purchased from Cell Signaling Technologies and used at 1:1000 dilution: cyclin D1 (2922S), Erk1/2 (9102S), phopho-Erk1/2 (9106S), merlin (12888S), phospho-merlin (13282S), GAPDH (5174S), and cyclophilin B (43603S). The vinculin antibody was purchased from Santa Cruz Biotechnologies (sc-25336) and used at 1:1000 dilution. The secondary antibodies were goat anti-rabbit (ThermoFisher Scientific, G-21234) and goat anti-mouse (ThermoFisher Scientific, G-21040). The secondary antibodies were used at 1:10,000 dilution.

### Microscopy

#### Brightfield imaging

Brightfield images were acquired on an inverted Nikon eclipse TS100 microscope using a 10X objective (Nikon, NA 1.2 Ph1 ADL) and a CCD camera (Q Imaging Retiga 01-RET-R3-R-M-14-C)

#### Epifluorescence imaging of labeled EdU and BAMPS in 2D

To image the labeled EdU and DAPI-stained nuclei, and for fast time-lapse 2D imaging of cells expressing mRuby2-tractin, we used an inverted epifluorescence Nikon Ti-Eclipse microscope equipped with a sCMOS C11449-22CU camera (Hamamatsu), SOLA solid state white-light excitation system, and a motorized filter turret with filters for DAPI, FITC, TRITC, and Cy5. Images were acquired using the Nikon Elements software. EdU and DAPI were imaged with 10X objective (Nikon NA 0.3 Ph1 DL). Dynamic BAMPs were imaged live using an oil-immersion 60X objective (Nikon NA 1.4 Plan Apo λ). Single-plane images were acquired every 2 seconds for 2 minutes. Cells imaged live were kept in a humidified chamber heated at 37°C at 5% CO^2^.

#### Light-sheet imaging

For volumetric imaging of cells cultured on 2D fibronectin-coated coverslips or in 3D collagen matrices, we used the Field Synthesis Gaussian light sheet microscope^21^, using a 25X long working distance and water immersion objective (Nikon NA 1.1, MRD77220). Samples were mounted on custom, 3D-printed holders. Cells were imaged in chambers filled with their appropriate growth media and heated to 37°C. Stacks were acquired at 300nm steps. The light intensity, exposure time, time interval, as well as the total acquisition time were optimized from sample to sample based on the brightness and the bleaching of the fluorophores.

#### Cleared-tissue light sheet imaging

We imaged cleared tissues using the cleared-tissue axially swept light-sheet microscope (ctASLM)^22^. We used a multi-immersion 36X objective (Advanced Scientific Imaging, NA 0.7) and sCMOS camera (Flash 4.0, Hamamatsu). Samples were imaged in CUBIC, the same solution used to clear the tissue. Before imaging, we incubated the samples in the imaging chamber in CUBIC overlayed with mineral oil overnight to equilibrate the refractive index. We imaged tissue sections of 0.5-2mm in thickness at a 200nm z step size. The light intensity and exposure time were optimized based on the brightness of the fluorophores.

#### Confocal imaging of the split SNAP-tag

Cells expressing the split SNAP-tag reporter were seeded on collagen-coated glass-bottom 35mm dishes (MatTek). For cells expressing myr-TIAM1DH/PH under the tetON promotor, doxycycline (2µg/mL) was added to the culture as cells were seeded. The cells were imaged on a Nikon SoRa Ti2 inverted spinning disk confocal microscope equipped with a Yokogawa CSU-W1 spinning disk unit, a Hamamatsu orca fusion BT CMOS digital camera and a 60X oil objective (Nikon, NA 0.85, Plan Apo λ Ph3 DM). Cells were imaged live, and a single stack was captured at 400nm z steps, such that the entire volume of the cell was covered.

#### Imaging of Proximity Ligation Assays (PLA)

We detected the PLA fluorescence signal in SU.86.86 cells using a Delta Vision OMX SR microscope system equipped with a 60X TIRF oil objective (Olympus, ApoN NA 1.49) and a scientific CMOS camera. The TIRF evanescent field depth was between 100 and 200nm. To capture the total number of spots in the cell, we imaged the whole cell volume using the same microscope and objective to acquire a z stack in the conventional acquisition mode, at a z step of 200nm.

#### Optogenetics

We used the MV3 cells for the optogenetics experiment. For light-induced membrane recruitment of the TIAM1 DH/PH domain, we used the Nikon SoRa Ti2 inverted spinning disk confocal microscope (see above). The iLID optogenetic module was triggered using the microscope’s 488nm laser exposed at 50ms at 50% laser power. Under these conditions, cells were imaged for 5 minutes. To increase the coverage of the light response and minimize user bias, we activated the entire field of view (as opposed to regions of interest on selected cells). To ensure that the observed TIAM1 DH/PH recruitment and subsequent BAMP formation were due to light activation, we imaged TIAM1DH/PH-tagRFP for 2 minutes before and for 5 minutes after 488nm laser activation. For the acquisition before, during, and after optogenetic activation, a single Z plane was acquired at a time interval of 2 seconds.

### Measuring the total cell fluorescence of the recovered SNAP-tag

We measured total cell fluorescence using ImageJ on sum projections of z stacks. On a sum projection, individual cells were carefully outlined using the free-hand selection tool on mRuby2 tractin images. Within the outlined cellular boundary, we measured the integrated density as the total cell fluorescence.

### Quantifying BAMP formation rate

We developed a ruffling rate quantification based on actin fluorescence live-cell imaging data. The ruffling rates were calibrated per video, where the field-of-view frame (FOV) typically contained 1-10 cells. Thresholding the areas with high actin intensities could not segment ruffles effectively, because of other actin-high structures such as stress fibers contaminated the readout. Instead, we exploited the fact that ruffles are more distinguishable in videos than static images because they move fast in space unlike stress fibers. This indicated that ruffling was represented as spikes in pixel intensity time courses. At each pixel and timepoint, we computed moving median intensities over a rolling time window of ±50 seconds (±25 frames) to extract low-frequency actin images. We then computed the ratio of the raw actin intensities to these images on a percentage scale to visualize regions with high frequency change in the actin signal. the regions with high normalized intensities were then delineated by the multi-scale-automatic (MSA) segmentation algorithm previously described^89^. The MSA algorithm identified ruffling areas in each time point via locally adaptive thresholding. The ruffling area was then related to the area of the entire cell, segmented separately in the raw actin image, to compute a percentage score of ruffling areas at each time point. Percentage scores were then averaged over time to define the ruffling rate of a live-cell video.

### Computation of SNAP-tag-KRASG12V subcellular enrichment

We first generated normalized SNAP-tag-KRASG12V movies by dividing each frame of the KRAS channel by the corresponding frame of the membrane marker channel. For each normalized movie, analysis depended on three user-defined parameters: 1) The intensity threshold for segmentation, which was manually tuned to allow separation of individual cells; 2) the first acquisition time point during light activation (T0), and 3) the final time point (T1). In the MV3 cellular model used in this experiment, the formation of BAMPs in response to light varied from cell to cell, but all responsive cells maintained appreciable BAMPs throughout light activation. Nonetheless, we noticed that upon prolonged light activation, BAMP formation decreased in some cells (See the Optogenetics section for light activation and image acquisition details). Therefore, for every cell analyzed, T1 was defined as the time point at which the cell reached maximum BAMP formation, as assessed by the user. The activating light was turned on every two seconds for 500 seconds, and for all the cells analyzed, T1 varied between 96 and 500 seconds. Each movie frame was foreground-background segmented using the determined threshold intensity. Protrusive parts of the cells were defined as those pixels which belonged to the cell (i.e. nonzero values) at T1 but not at T0. Pixels that belonged to the cells at both T0 and T1 were considered to be outside the protrusions (non-protrusive). To quantify subcellular enrichment of KRAS in BAMPs, we computed and compared the mean normalized SNAP-tag-KRASG12V intensity inside and outside protrusions.

### Proximity ligation assay

We used Duolink far red detection reagents from Millipore Sigma (DUO92013-30RXN). The anti-mouse MINUS probe (DUO92004-30RXN), the anti-rabbit PLUS probe (DUO92002- 30RXN), and the wash buffers (DUO82049-4L) were also purchased from Millipore Sigma. Cells were cultured on collagen-coated glass-bottom 35mm dishes with a 7mm coverslip (MatTek, P35G-1.5-7-C) and incubated overnight. For the expression of myr-TIAM1DH/PH under the tetON promotor, doxycycline (2µg/mL) was added to the culture as the cells were seeded. We fixed the cells with 4% paraformaldehyde in PBS for 10 minutes at 37°C. Cells were permeabilized for 20 minutes at room temperature using 0.5% triton in 5% donkey serum in PBS. After washing 3 times with PBS, we completed the PLA according to the manufacturer’s instructions. Briefly, blocking was done for 1 hour at 37°C before primary antibody incubation overnight at 4°C. We then incubated the cells with the PLA probes for 1 hour at 37°C. Ligation was done for 30 minutes at 37°C, and the amplification reaction was incubated for 100 minutes at 37°C. All the washes between steps were done as instructed by the manufacturer, and all the incubations were done in a humidified chamber. After the PLA, cells were kept at 4°C in PBS until imaging. Cells were imaged in PBS.

### PLA spot detection and counting

In TIRF images, the PLA spots were counted manually. To distinguish the lamellipodia from the rest of the cell in SU.86.86, we carefully traced the outer boundary of the lamella region, which was marked by pronounced linear actin filaments, using the free-hand tool in ImageJ. To calculate the spot density in each cellular compartment, we divided the number of PLA spots within the compartment by its area.

To detect and count all the spots in Z stack acquisitions of MIA-PaCa-2 cells, the input 3D image was resized to be isotropic. Intensities were then percentile normalized to be 0-1 where 0 and 1 correspond to the 2^nd^ and 99.8^th^ percentile intensity values in the original. Spots were then detected using the difference of Gaussian method using the blob dog function in Python Scikit-image library (skimage.feature.blob_dog with parameters, min_sigma=3, max_sigma=6, threshold=0.01). Lastly, we eliminated spurious spot detections outside the cell volume by deriving from the actin channel image a binary cell mask based on 3-class Otsu thresholding to keep the brightest voxels. This initial segmentation was postprocessed to keep the largest spatial component and performing binary morphological closing (ball kernel radius 3), binary filling holes, and binary morphological erosion (ball kernel radius 1).

### Cell image segmentation

The 3D images were preprocessed by gamma correction (gamma=0.8) and median filtering (neighborhood size = 3 x 3 x 3 voxels). The image was then interpolated to be isotropic, and its intensities percentile normalized to be 0-1 where 0 and 1 correspond to the 2^nd^ and 99.8^th^ percentile intensity values in the original. This image was downsampled isotropically by 1/4 in each axis to enable a complete cell segmentation. Binary Otsu thresholding was performed to segment each 2D slice in x-y, x-z, and y-z orthoviews with a minimum normalized intensity of 0.2. Then, the 2D segmented stacks from each view were combined into a single consensus 3D segmentation using u-Segment3D^90^, and retaining the largest spatial component. This 3D segmentation was further postprocessed as described in u-Segment3D to recover fine surface protrusions, first by label diffusion using the segmentation input image as the guide image, and second by resizing the segmentation to the original isotropic resolution to perform guided filtering with a ridge feature enhanced version of the original 3D image as the guide image.

### Proximal surface fluorescence signal mapping

The surface proximal signal intensities of Arp2/3, Tractin and Ras were mapped using u-Unwrap3D^91^. The procedure averaged the intensities along a curvilinear trajectory of 1µm length from the segmented cell surface along the gradient of the Euclidean distance transform of the cell volume into the cell interior. The surface intensities were rendered on the mesh using the RdYlBu colormap with the minimum and maximum color range set to the 1^st^ and 99^th^ percentile of the intensities.

## Data availability and code availability

Data supporting the presented findings can be found here: https://doi.org/10.5281/zenodo.19224721

Computer codes used in this study are publicly available online:

Ruffling rate quantification: DOI 10.5281/zenodo.18763493 and https://github.com/JungsikNoh/RuffleSegmentation

3D cell image segmentation: https://github.com/DanuserLab/u-segment3D

3D Spot detection (for PLA data): https://scikit-image.org/docs/stable/auto_examples/features_detection/plot_blob.html

Proximal cell surface fluorescence signal mapping: https://github.com/DanuserLab/u-unwrap3D

Computation of oncogenic KRAS enrichment in cell membrane protrusions: https://github.com/DanuserLab/Gihana_BAMPS2026

## Acknowledgements

We thank M. Metlen, D. Mundy, and Etai Sapoznik for their guidance in microscopy experiments. We thank A. Weems for discussions on interventional approaches to inhibit BAMP formation. We thank Qiongjing Zou for the her support in curating and publishing computer code, and we thank D. Reed for logistical support throughout the study.

## Author Contributions

G. Gihana conceived and coordinated the project, designed and performed biochemical and imaging experiments, and analyzed data. G. Gihana and G. Danuser wrote the manuscript. G. Danuser, K. Dean, M. Conacci-Sorrell, and R. Fiolka guided the conception and the development of the study, and they secured funding. G. Danuser supervised the study as the principal investigator. K. Bhatt designed and performed western blot and CRISPR experiments. B. Chang designed and performed light sheet imaging experiments and processed the images. M. Vasanth performed cell proliferation assays. F. Zhou computed cell surface fluorescence signal mapping and PLA spot detection. J. Noh computed the membrane ruffling rate. R. Ravishankar computed oncogenic KRAS subcellular enrichment. H. Borges cleared tumor tissues. J. Lin optimized light sheet imaging of cleared tissues. PD Nogueira, L. Perez Castro, and N. Venkateswaran performed mouse xenografts. B. Chen optimized and performed light sheet imaging of cell spheroids.

## Corresponding authors

Gaudenz Danuser and Gabriel Muhire Gihana

## Electronic Supplementary Material

**Supplementary Video 1: BAMP formation in 2D culture of SU.86.86 cells.**

Maximum intensity projection images of an SU.86.86 cell expressing both CyOFP1-tractin and Arp3-EGFP. The cells were plated on a fibronectin-coated coverslip overnight and imaged live with a gaussian light sheet on the Field Synthesis microscope. Images were acquired every 11 seconds and animated at seven frames per second. Scale bar: 20µm.

**Supplementary Video 2: BAMP formation in 3D culture of SU.86.86 cells**

Maximum intensity projection images of an SU.86.86 cell expressing both CyOFP1-tractin and Arp3-EGFP. The cells were cultured in 3D collagen matrix overnight and imaged live using a gaussian light sheet on the Field Synthesis microscope. Images were acquired every 8 seconds and animated at seven frames per second. Scale bar: 20µm.

**Supplementary Video 3: SU.86.86 spheroids**

Maximum intensity projection images of SU.86.86 cells expressing mEmerald-tractin. The video shows a cell spheroid formed after two weeks of incubation in 3D collagen culture. The cells were imaged using a gaussian light sheet on an oblique-plane light sheet microscope (OPM)^23^. BAMPs appear as dynamic, F-actin-rich cortical protrusions. Images were acquired every 9 seconds and animated at seven frames per second. Scale bar: 30µm.

**Supplementary Video 4: Animated maximum intensity projection of SU.86.86 cells in a mouse subcutaneous xenograft**

The representative cells were selected from tissue sections of a subcutaneous mouse xenograft tumor. Cells express CyOFP1-tractin. For representation of BAMPs, we selected cells localized at the periphery of the tumor mass.

**Supplementary Video 5: BAMP formation in 2D culture of COR-L23 cells.**

Maximum intensity projection images of a COR-L23 cell expressing both mRuby2-tractin. The cells were plated on a fibronectin-coated coverslip overnight and imaged live on the Field Synthesis gaussian light sheet microscope. Scale bar: 10µm.

**Supplementary Video 6: BAMP formation in 3D culture of COR-L23 cells**

Maximum intensity projection images of a COR-L23 cell expressing both mRuby2-tractin. The cells were cultured in 3D collagen matrix overnight and imaged live on the Field Synthesis gaussian light sheet microscope. Scale bar: 10µm.

**Supplementary Video 7: Animated maximum intensity projection of COR-L23 cells in a mouse subcutaneous xenograft**

The representative cells were selected from tissue sections of a subcutaneous mouse xenograft tumor of COR-L23 cells expressing CyOFP1-tractin. For representation of BAMPs, we selected cells localized at the periphery of the tumor mass.

---

**Extended Data Fig.1:**
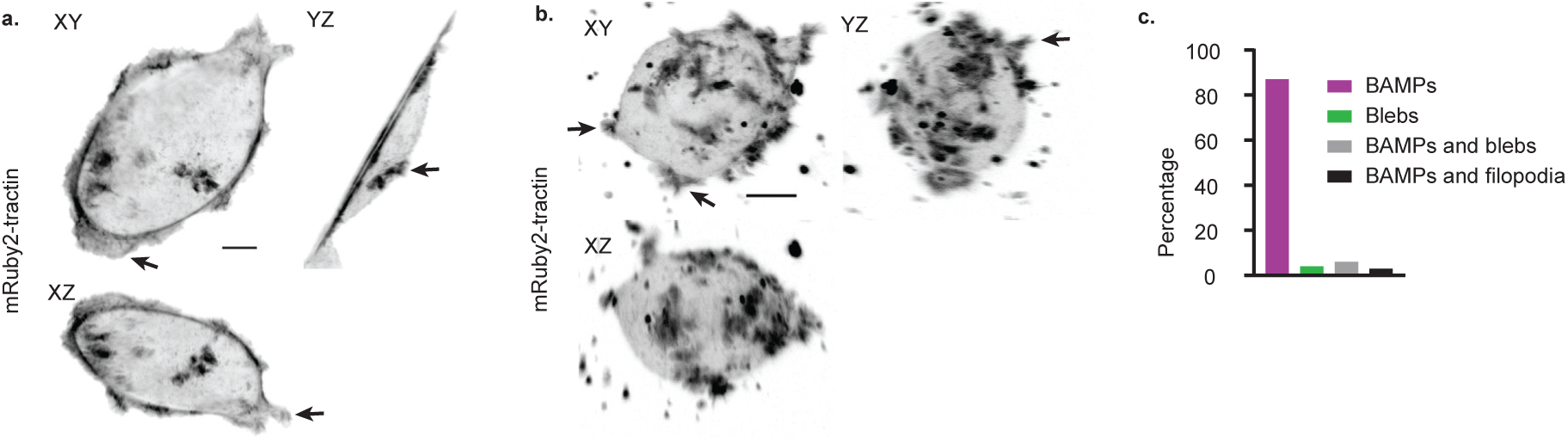
BAMP formation in COR-L23 human lung carcinoma cells. **(a)** Maximum intensity projection (MIP) images of a representative COR-L23 cell in 2D culture. The cells were plated on a fibronectin-coated coverslip overnight before imaging. Arrows in this and panel b) indicate examples of BAMPs extending from the cortical surface of the cell. Scale bar: 10µm. **(b)** MIP images of a representative COR-L23 cell in 3D collagen culture. The cells were cultured overnight before imaging. Scale bar: 10µm. **(c)** Frequency of plasma membrane protrusion types formed by COR-L23 cells cultured in 3D collagen matrix. Images were acquired with a gaussian light sheet microscope.

**Extended Data Fig.2:**
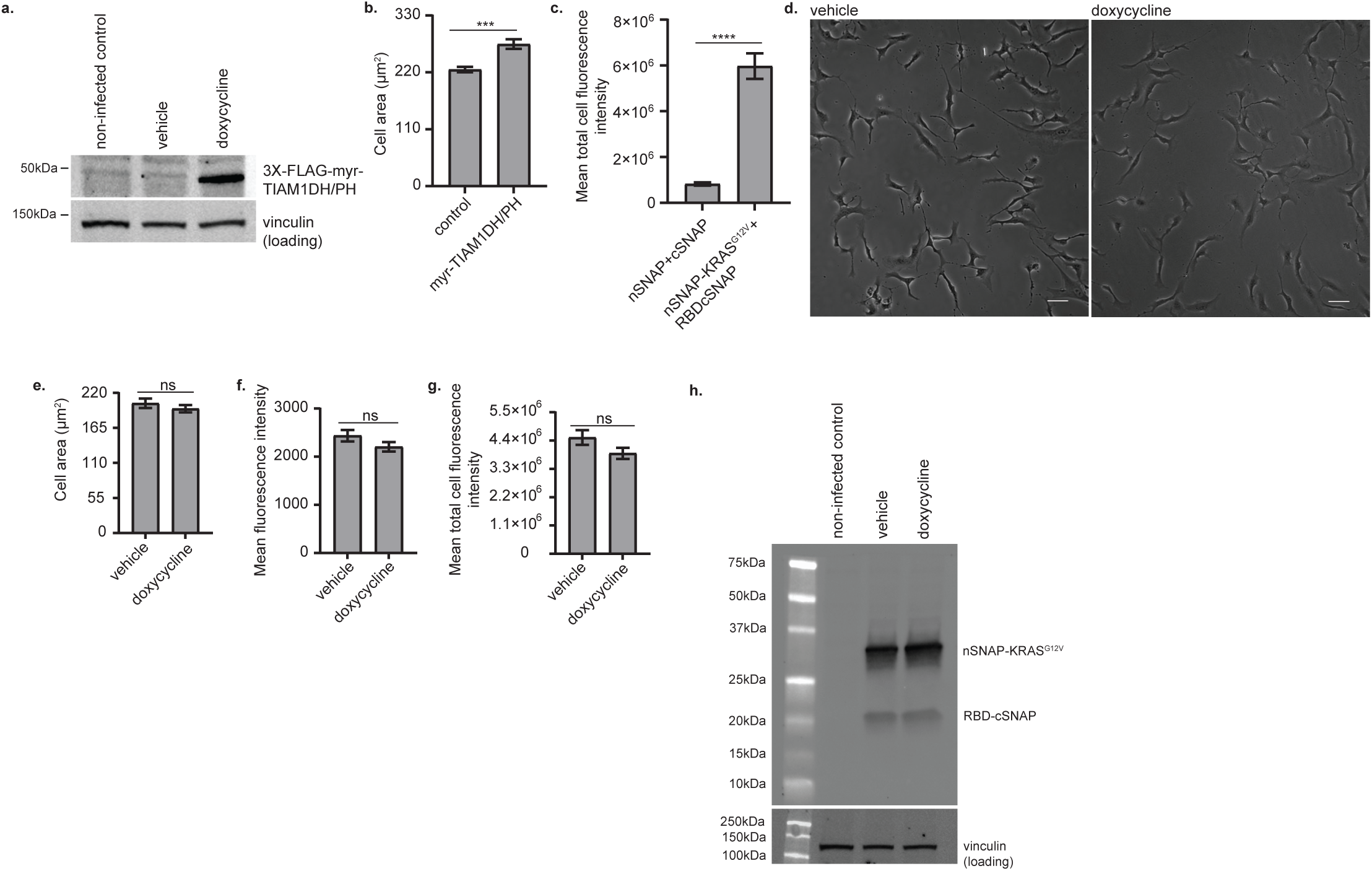
Split SNAP-tag controls. **(a)** Doxycycline-induced expression of the FLAG-tagged myr-TIAM1DH/PH in HPNE cells. Non-infected control: HPNE cells that did not carry the plasmid harboring the 3XFLAG-myr-TIAM1DH/PH. Vehicle: water solvent. Vinculin was used as a loading control. The expected size of 3XFLAG-myr-TIAM1DH/PH is 48.2kDa. **(b)** Area of HPNE cells containing doxycycline inducible myr-TIAM1DH/PH and treated with the water solvent vehicle (control) or doxycycline (myr-TIAM1DH/PH). (n = 224 cells for each condition from three independent replicates; ***: p value = 0.0001). **(c)** Split SNAP-tag total cell fluorescence intensity in HPNE cells expressing doxycycline-induced myr-TIAM1DH/PH. (n = 73 cells from one replicate; ****: p value < 0.0001). **(d)** Brightfield images of HPNE cells expressing nSNAP-KRASG12V and RBD-cSNAP fragments of the Split SNAP-tag and treated with either water(vehicle) or doxycycline. Scale bar: 40µm. **(e)** Area of HPNE cells expressing both nSNAP-KRASG12V and RBD-cSNAP and treated with either the vehicle or doxycycline. **(f, g)** Split SNAP-tag mean and total fluorescence intensity, respectively, in HPNE cells cells expressing both nSNAP-KRASG12V and RBD-cSNAP. For e, f, g, the cells did not contain myr-TIAM1DH/PH, and all the plots show the mean with SEM error bars. P values were calculated using a two-tailed Mann-Whitney test (n = 101 from one replicate; ns: p value > 0.05). **(h)** Expression of nSNAP-KRASG12V and RBD-cSNAP fragments of the Split SNAP-tag in HPNE cells infected with doxycycline-inducible and FLAG-tagged myr-TIAM1DH/PH. The non-infected control cells did not carry any integrated plasmid. Vehicle: water solvent. Expected sizes for nSNAP-KRASG12V and RBD-cSNAP are 31.4kDa and 20.2kDa, respectively.

**Extended Data Fig.3:**
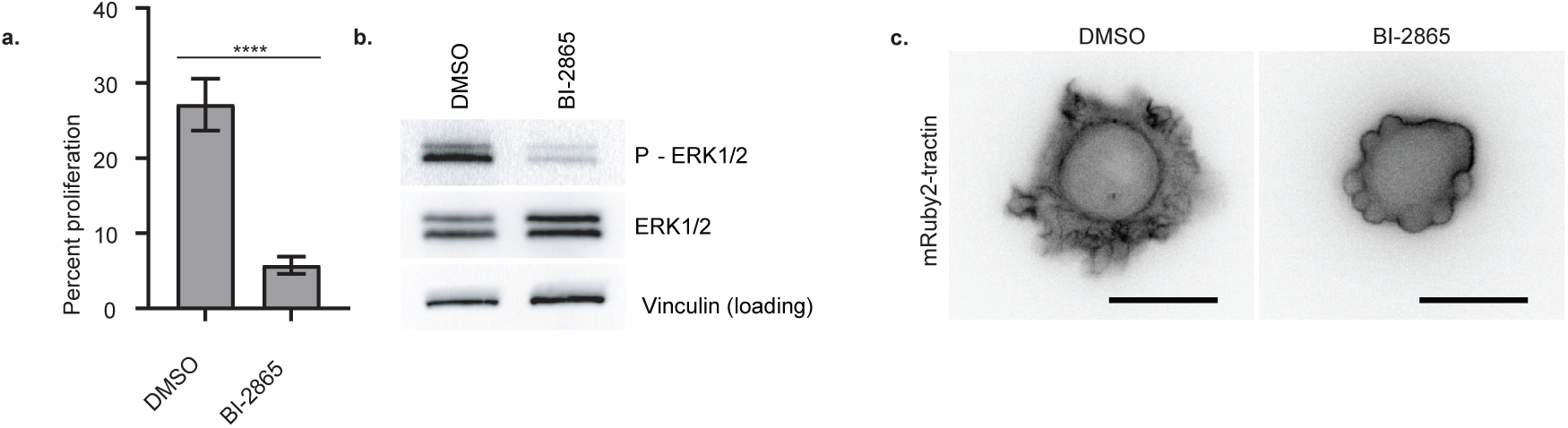
KRAS inhibition attenuates BAMPs in SU.86.86 cells. **(a)** Proliferation rate of SU.86.86 cells treated with DMSO or BI-28655. The plot shows the mean with SEM error bars (n = 30 images from 3 independent replicates, with each image containing dozens of cells; ****: p value <0.0001, two-tailed Mann-Whitney test). **(b)** Western blotting for phosphorylated ERK1/2 in SU.86.86 cells treated with DMSO or BI-2865. Vinculin was used as a loading control. **(c)** Epifluorescence images of SU.86.86 cells treated with DMSO or BI-2865. Scale bar: 20µm.

**Extended Data Fig.4:**
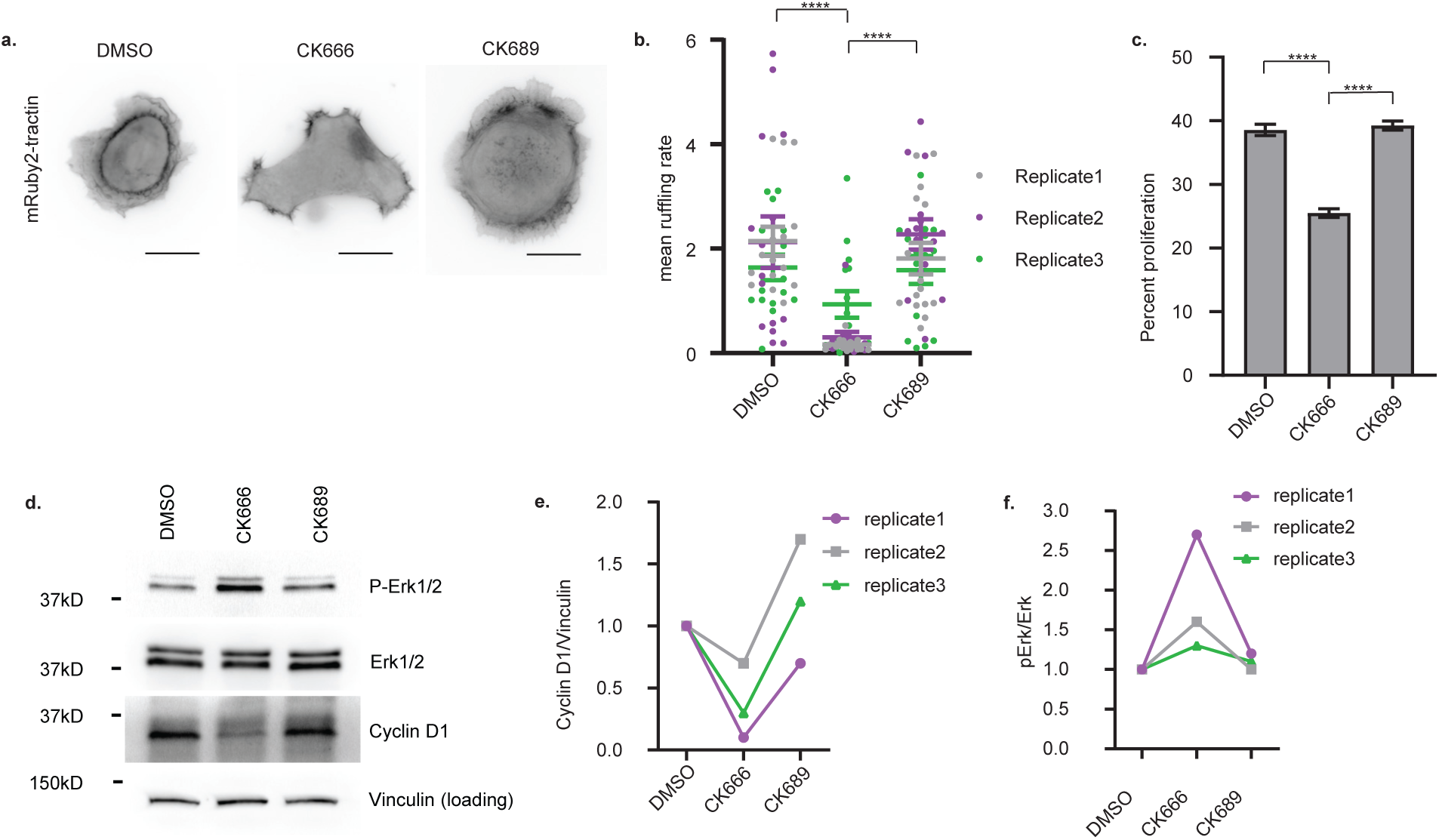
BAMPs promote cyclin D1 expression and cell proliferation downstream of oncogenic KRAS, independently of MAPK in COR-L23 Cells. **(a)** Epifluorescence images of COR-L23 cells treated with DMSO, CK666, or CK689. Scale bar: 10µm. **(b)** BAMP formation quantified as the mean ruffling rate computed in the cells represented in (a). Dots represent individual images; error bars represent SEM (n = 42 images from 3 independent replicates, with each image containing 1-10 cells; ****: p value <0.0001, two-tailed Mann-Whitney test). **(c)** Proliferation rate of COR-L23 cells treated with DMSO, CK666, or CK689. The plot shows the mean with SEM error bars (n = 38 images from three independent replicates, with each image containing dozens of cells; ****: p value <0.0001, two-tailed Mann-Whitney test). **(d)** Western blotting for phosphorylated ERK1/2 and cyclin D1 in COR-L23 cells treated DMSO, CK666, or CK689. **(e, f)** Quantification of band intensity on the western blot images represented in (d). DMSO was used at 0.2%, CK666 and CK689 at 200µM.

