## Supplementary Table 1 for "Modulation of Oncogenic KRAS Signaling by Branched Actin-driven Cell Membrane Protrusions"

Plasmid List

| Plasmid | Source |
|---|---|
| pLVX_IRES_Neomycin empty vector | Wesley Burford (UTSW) |
| pLVX_IRES_Puromycin empty vector | Wesley Burford (UTSW) |
| pTRE3G empty vector | This study |
| pLVX:mRuby2-tractin | Wesley Burford (UTSW) |
| pLVX:CyOFP1-tractin | Wesley Burford (UTSW) |
| pLVX:mEmerald-tractin | Wesley Burford (UTSW) |
| pLVX:Arp3-EGFP | Tadamoto Isogai (UTSW) |
| pLVX:EGFP-KRASG12V | This study |
| pLVX:SNAP-tag-KRASG12V | This study |
| pLVX:nSNAP-KRASG12V | This study |
| pLVX:RBD-cSNAP | This study |
| pLL7.0: mTiam1(64-437)-tgRFPt-SSPB R73Q | Addgene #60418 |
| pLL7.0: Venus-iLID-CAAX | Addgene #60411 |
| pLVX:EGFP-CAAX | This study |
| pTRE3G:myr-TIAM1DH/PH | This study |
| pTRE3G:NF2WT | This study |
| pTRE3G:NF2S518A | This study |
| pMD.2g | Addgene #12259 |
| psPax2 | Addgene #12260 |
| pSpCas9(BB)-2A-Puro | Addgene # 62988 |
| pMA-Tia1L | Tilmann Bürckstümmer (Horizon Genomics) |

nSNAP: Amino terminus (N-terminus) fragment of the SNAP-tag protein (amino acids 1-91)

cSNAP: Carboxyl terminus (C-terminus) fragment of the SNAP-tag protein (amino acids 92-182).

RBD: RAS Binding Domain of the c-RAF1 effector kinase (amin acids 51-131).
