## Extended Data File 1 for "Modulation of Oncogenic KRAS Signaling by Branched Actin-driven Cell Membrane Protrusions"

#### ENDOGENOUS KRAS SEQUENCE VERIFICATION

**COR-L23 Cell Line**

Seq_1: sequence of the Reference WT KRAS DNA (NCBI reference: NM_004985.5)
Seq_2: sequence of the product of a polymerase chain reaction (PCR) targeted at the KRAS endogenous locus

```
Seq_1  1      aagcgtcgatggaggagtttgtaaatgaagtacagttcattacgatacacgtctgcagtc  60
                           |||||||||||||||||||||||||||||||||||||||||||||||
Seq_2  1      -------------ggagtttgtaaatgaagtacagttcattacgatacacgtctgcagtc  47


Seq_1  61     aactggaattttcatgattgaattttgtaaggtattttgaaataatttttcatataaagg  120
              ||||||||||||||||||||||||||||||||||||||||||||||||||||||||||||
Seq_2  48     aactggaattttcatgattgaattttgtaaggtattttgaaataatttttcatataaagg  107


Seq_1  121    tgagtttgtattaaaaggtactggtggagtatttgatagtgtattaaccttatgtgtgac  180
              ||||||||||||||||||||||||||||||||||||||||||||||||||||||||||||
Seq_2  108    tgagtttgtattaaaaggtactggtggagtatttgatagtgtattaaccttatgtgtgac  167

                                                                   M  T  E
Seq_1  181    atgttctaatatagtcacattttcattatttttattataagGCCTGCTGAAAATGACTGA  240
              ||||||||||||||||||||||||||||||||||||||||||||||||||||||||||||
Seq_2  168    atgttctaatatagtcacattttcattatttttattataaggcctgctgaaaATGACTGA  227
                                                                   M  T  E

               Y  K  L  V  V  V  G  A  G  G  V  G  K  S  A  L  T  I  Q  L
Seq_1  241    ATATAAACTTGTGGTAGTTGGAGCTGGTGGCGTAGGCAAGAGTGCCTTGACGATACAGCT  300
              |||||||||||||||||||||||||| |||||||||||||||||||||||||||||||||
Seq_2  228    ATATAAACTTGTGGTAGTTGGAGCTGTTGGCGTAGGCAAGAGTGCCTTGACGATACAGCT  287
               Y  K  L  V  V  V  G  A  V  G  V  G  K  S  A  L  T  I  Q  L

               I  Q  N  H  F  V  D  E  Y  D  P  T  I  E
Seq_1  301    AATTCAGAATCATTTTGTGGACGAATATGATCCAACAATAGAGgtaaatcttgttttaat  360
              ||||||||||||||||||||||||||||||||||||||||||||||||||||||||||||
Seq_2  288    AATTCAGAATCATTTTGTGGACGAATATGATCCAACAATAGAGgtaaatcttgttttaat  347
               I  Q  N  H  F  V  D  E  Y  D  P  T  I  E

Seq_1  361    atgcatattactggtgcaggaccattctttgatacagataaaggtttctctgaccatttt  420
              ||||||||||||||||||||||||||||||||||||||||||||||||||||||||||||
Seq_2  348    atgcatattactggtgcaggaccattctttgatacagataaaggtttctctgaccatttt  407


Seq_1  421    catgagtacttattacaagataattatgctgaaagttaagttatctgaaatgtaccttgg  480
              |||
Seq_2  408    cat---------------------------------------------------------  410
```

Extended Data File 1

**SU.86.86 Cell Line**

Seq_1: sequence of the Reference WT KRAS DNA (NCBI reference: NM_004985.5)
Seq_2: sequence of the product of a polymerase chain reaction (PCR) targeted at the KRAS endogenous locus

```
Seq_1  1      aagcgtcgatggaggagtttgtaaatgaagtacagttcattacgatacacgtctgcagtc  60
                           ||||||||||||||||||||||||||||||||||||||||||||||
Seq_2  1      -------------ggagtttgtaaatgaagtacagttcattacgatacacgtctgcagtc  47


Seq_1  61     aactggaattttcatgattgaattttgtaaggtattttgaaataatttttcatataaagg  120
              ||||||||||||||||||||||||||||||||||||||||||||||||||||||||||||
Seq_2  48     aactggaattttcatgattgaattttgtaaggtattttgaaataatttttcatataaagg  107


Seq_1  121    tgagtttgtattaaaaggtactggtggagtatttgatagtgtattaaccttatgtgtgac  180
              ||||||||||||||||||||||||||||||||||||||||||||||||||||||||||||
Seq_2  108    tgagtttgtattaaaaggtactggtggagtatttgatagtgtattaaccttatgtgtgac  167

                                                                  M  T  E
Seq_1  181    atgttctaatatagtcacattttcattatttttattataagGCCTGCTGAAAATGACTGA  240
              ||||||||||||||||||||||||||||||||||||||||||||||||||||||||||||
Seq_2  168    atgttctaatatagtcacattttcattatttttattataagGCCTGCTGAAAATGACTGA  227
                                                                  M  T  E

               Y  K  L  V  V  V  G  A  G  G  V  G  K  S  A  L  T  I  Q  L
Seq_1  241    ATATAAACTTGTGGTAGTTGGAGCTGGTGGCGTAGGCAAGAGTGCCTTGACGATACAGCT  300
              ||||||||||||||||||||||||| ||||||||||||||||||||||||||||||||||
Seq_2  228    ATATAAACTTGTGGTAGTTGGAGCTGATGGCGTAGGCAAGAGTGCCTTGACGATACAGCT  287
               Y  K  L  V  V  V  G  A  D  G  V  G  K  S  A  L  T  I  Q  L

               I  Q  N  H  F  V  D  E  Y  D  P  T  I  E
Seq_1  301    AATTCAGAATCATTTTGTGGACGAATATGATCCAACAATAGAGgtaaatcttgttttaat  360
              ||||||||||||||||||||||||||||||||||||||||||||||||||||||||||||
Seq_2  288    AATTCAGAATCATTTTGTGGACGAATATGATCCAACAATAGAGgtaaatcttgttttaat  347
               I  Q  N  H  F  V  D  E  Y  D  P  T  I  E

Seq_1  361    atgcatattactggtgcaggaccattctttgatacagataaaggtttctctgaccatttt  420
              ||||||||||||||||||||||||||||||||||||||||||||||||||||||||||||
Seq_2  348    atgcatattactggtgcaggaccattctttgatacagataaaggtttctctgaccatttt  407


Seq_1  421    catgagtacttattacaagataattatgctgaaagttaagttatctgaaatgtaccttgg  480
              |||||||||
Seq_2  408    catgagtac---------------------------------------------------  416
```

### Extended Data File 1

**MIA PaCa-2 Cell Line**

Seq_1: sequence of the Reference WT KRAS DNA (NCBI reference: NM_004985.5)
Seq_2: sequence of the product of a polymerase chain reaction (PCR) targeted at the KRAS endogenous locus

```
Seq_1  1     aagcgtcgatggaggagtttgtaaatgaagtacagttcattacgatacacgtctgcagtc  60
                         ||||||||||||||||||||||||||||||||||||||||||||||||
Seq_2  378   ------------aggagtttgtaaatgaagtacagttcattacgatacacgtctgcagtc  331


Seq_1  61    aactggaattttcatgattgaattttgtaaggtatttgaaataatttttcatataaagg  120
             |||||||||||||||||||||||||||||||||||||||||||||||||||||||||||
Seq_2  330   aactggaattttcatgattgaattttgtaaggtatttgaaataatttttcatataaagg  271


Seq_1  121   tgagtttgtattaaaaggtactggtggagtatttgatagtgtattaaccttatgtgtgac  180
             ||||||||||||||||||||||||||||||||||||||||||||||||||||||||||||
Seq_2  270   tgagtttgtattaaaaggtactggtggagtatttgatagtgtattaaccttatgtgtgac  211


                                                                M  T  E
Seq_1  181   atgttctaatatagtcacattttcattatttttattataagGCCTGCTGAAAATGACTGA  240
             ||||||||||||||||||||||||||||||||||||||||||||||||||||||||||||
Seq_2  210   atgttctaatatagtcacattttcattatttttattataaggcctgctgaaaATGACTGA  151
                                                                M  T  E


              Y  K  L  V  V  V  G  A  G  G  V  G  K  S  A  L  T  I  Q  L
Seq_1  241   ATATAAACTTGTGGTAGTTGGAGCTGGTGGCGTAGGCAAGAGTGCCTTGACGATACAGCT  300
             ||||||||||||||||||||||||| |||||||||||||||||||||||||||||||||||
Seq_2  150   ATATAAACTTGTGGTAGTTGGAGCTTGTGGCGTAGGCAAGAGTGCCTTGACGATACAGCT  91
              Y  K  L  V  V  V  G  A  C  G  V  G  K  S  A  L  T  I  Q  L


              I  Q  N  H  F  V  D  E  Y  D  P  T  I  E
Seq_1  301   AATTCAGAATCATTTTGTGGACGAATATGATCCAACAATAGAGgtaaatcttgttttaat  360
             ||||||||||||||||||||||||||||||||||||||||||||||||||||||||||||
Seq_2  90    AATTCAGAATCATTTTGTGGACGAATATGATCCAACAATAGAGgtaaatcttgttttaat  31
              I  Q  N  H  F  V  D  E  Y  D  P  T  I  E


Seq_1  361   atgcatattactggtgcaggaccattctttgatacagataaaggtttctctgaccatttt  420
             ||||||||||||||||||||||||||||||
Seq_2  30    atgcatattactggtgcaggaccattcttt------------------------------  1


Seq_1  421   catgagtacttattacaagataattatgctgaaagttaagttatctgaaatgtaccttgg  480

Seq_2  0     ------------------------------------------------------------  1
```

**Extended Data File 1: Verification of KRAS mutation at codon 12 in patient-derived cell lines.** The PCR product sequence was aligned to a refence wildtype KRAS4B sequence (NCBI reference: NM_004985.5) to validate the KRAS mutations at codon 12. For every sequence, the first methionine codon is highlighted in green. Codon 12 is highlighted in blue and magenta for the reference and PCR sequences, respectively. Yellow highlights the amino acid at codon 12.
